# Estimating the functional dimensionality of neural representations

**DOI:** 10.1101/232454

**Authors:** Christiane Ahlheim, Bradley C. Love

## Abstract

Recent advances in multivariate fMRI analysis stress the importance of information inherent to voxel patterns. Key to interpreting these patterns is estimating the underlying dimensionality of neural representations. Dimensions may correspond to psychological dimensions, such as length and orientation, or involve other coding schemes. Unfortunately, the noise structure of fMRI data inflates dimensionality estimates and thus makes it difficult to assess the true underlying dimensionality of a pattern. To address this challenge, we developed a novel approach to identify brain regions that carry reliable task-modulated signal and to derive an estimate of the signal’s functional dimensionality. We combined singular value decomposition with cross-validation to find the best low-dimensional projection of a pattern of voxel-responses at a single-subject level. Goodness of the low-dimensional reconstruction is measured as Pearson correlation with a test set, which allows to test for significance of the low-dimensional reconstruction across participants. Using hierarchical Bayesian modeling, we derive the best estimate and associated uncertainty of underlying dimensionality across participants. We validated our method on simulated data of varying underlying dimensionality, showing that recovered dimensionalities match closely true dimensionalities. We then applied our method to three published fMRI data sets all involving processing of visual stimuli. The results highlight three possible applications of estimating the functional dimensionality of neural data. Firstly, it can aid evaluation of model-based analyses by revealing which areas express reliable, task-modulated signal that could be missed by specific models. Secondly, it can reveal functional differences across brain regions. Thirdly, knowing the functional dimensionality allows assessing task-related differences in the complexity of neural patterns.

## 1. Introduction

A growing number of fMRI studies are investigating the representational geometry of voxel response patterns. For example, using representational similarity analysis (RSA; Kriegeskorte and Kievit, 2013), researchers have characterized visual object representations along the ventral stream (Khaligh-Razavi and Kriegeskorte, 2014) and how these representations vary across tasks (Bracci et al., 2017).

Interpreting representational geometry in neural responses can be difficult. For example, RSA tests for a hypothesized representational pattern, but an important and more fundamental question should be addressed first, namely whether there is any dimensionality to the underlying neural pattern and, if so, what that dimensionality is.

Knowing whether a pattern has dimensionality should be prerequisite for RSA and other multivariate representational analyses because a particular similarity structure can only be found when there is sufficient dimensionality to represent the proposed relations. For example, searching for a flavor space with dimensions sweet, sour, bitter, salty and umami would be a fool’s errand in brain areas that contain little or no dimensionality.

Although previous studies have made substantial progress in identifying whether any dimensionality underlies an observed pattern (Naselaris et al., 2011; Diedrichsen et al., 2016; Walther et al., 2016; Allefeld and Haynes, 2014), a straightforward, general, robust, open source, and computationally efficient procedure for this challenge would be welcomed. Moreover, progress would be welcomed on perhaps the more challenging task of estimating the degree of dimensionality underlying a pattern. Independent of the particular geometry, the dimensionality of a neural pattern is informative of how many features of a task are represented in a brain region, which can inform our understanding of an area’s function.

There are many methods of dimensionality reduction and estimation, most of which involve low-rank matrix approximation and aim to maximize the correspondence between the original and the approximated matrix. For example, two common approaches to estimate the dimensionality of an observed neural or behavioral pattern are principal component analysis (PCA) or relatedly, multidimensional scaling (MDS).

PCA, or the closely related factor analysis and singular value decomposition (SVD) (Hastie et al., 2009), is widely used in the study of individual differences and aids estimating how many latent components, or “factors”, underlie a pattern of (item) responses within or across participants, as for instance in the context of intelligence (Spearman, 1904) or personality tests (Cattell, 1947). In the context of neuroimaging, PCA has been used to identify brain networks (Huth et al., 2012; Friston et al., 1993). PCA derives how much variance of the observed pattern is explained by each underlying component.

Similarly, MDS finds the best representation of original distances in a low-dimensional space (Kriegeskorte and Kievit, 2013). For example, two stimuli like a chair and table that are very close to each other in the high-dimensional space will be represented closely in the low-dimensional projection achieved by MDS, whereas two stimuli that were very distant from each other, for instance a chair and a bunny, will be projected far apart. MDS has been successfully applied to behavioral as well as neural data to reveal which stimulus features underly observed representational geometries (Bracci and Op de Beeck, 2016; Kriegeskorte and Kievit, 2013; Kriegeskorte et al., 2008), though it has been questioned to which extent results from MDS are interpretable (Goddard et al., 2017). For reasons outlined below, we will focus on SVD to estimate the dimensionality of neural representations, though other methods could be paired with our general approach, including nonlinear approaches such as Nonlinear PCA (Kramer, 1991).

Estimating the dimensionality of neural data brings its own unique challenges. In a noise-free scenario, dimensionality can be defined as the number of linear orthogonal components (singular- or eigenvalues) underlying a matrix that are larger than zero (Shlens, 2014), indicating that the component fits some variance in the data. Unfortunately, actual recordings of neural activity always contain noise, which inflates non-signal components above zero (Fusi et al., 2016; Diedrichsen et al., 2013). This noise makes it challenging to determine which areas contain signal and, if so, what the dimensionality of the signal is.

One criterion, which we adopt in the work reported here, is to choose the number of components that should maximize reconstruction accuracy (measured by correlation) on new data (i.e., test data). While even for data with low or moderate true dimensionality more components will always increase fit for existing data (i.e., training data), performance on test data (i.e., generalization, prediction) will usually be best for a moderate number of components because these components largely reflect true signal as opposed to noise in the observed training sample.

The problem of distinguishing between signal and noise in a neural pattern is related to the bias-variance trade-off in supervised learning and model-selection. Overly simple models (few components) are highly biased, fitting training data poorly and not performing well on test data. These overly simple models cannot pick-up on nuances in the signal. Conversely, overly complex models (many components) are too sensitive to the variance in the training date (i.e., overfit). Although they fit the training data very well, overly complex models treat noise in the training data as signal and, therefore, generalize poorly. Thus, the sweet spot for test performance should be at some moderate number of components that largely reflect true signal (see Figure 1 A). Thus, identifying the true number of underlying components is analogous to deciding which model best explains the data.

**Figure 1:**
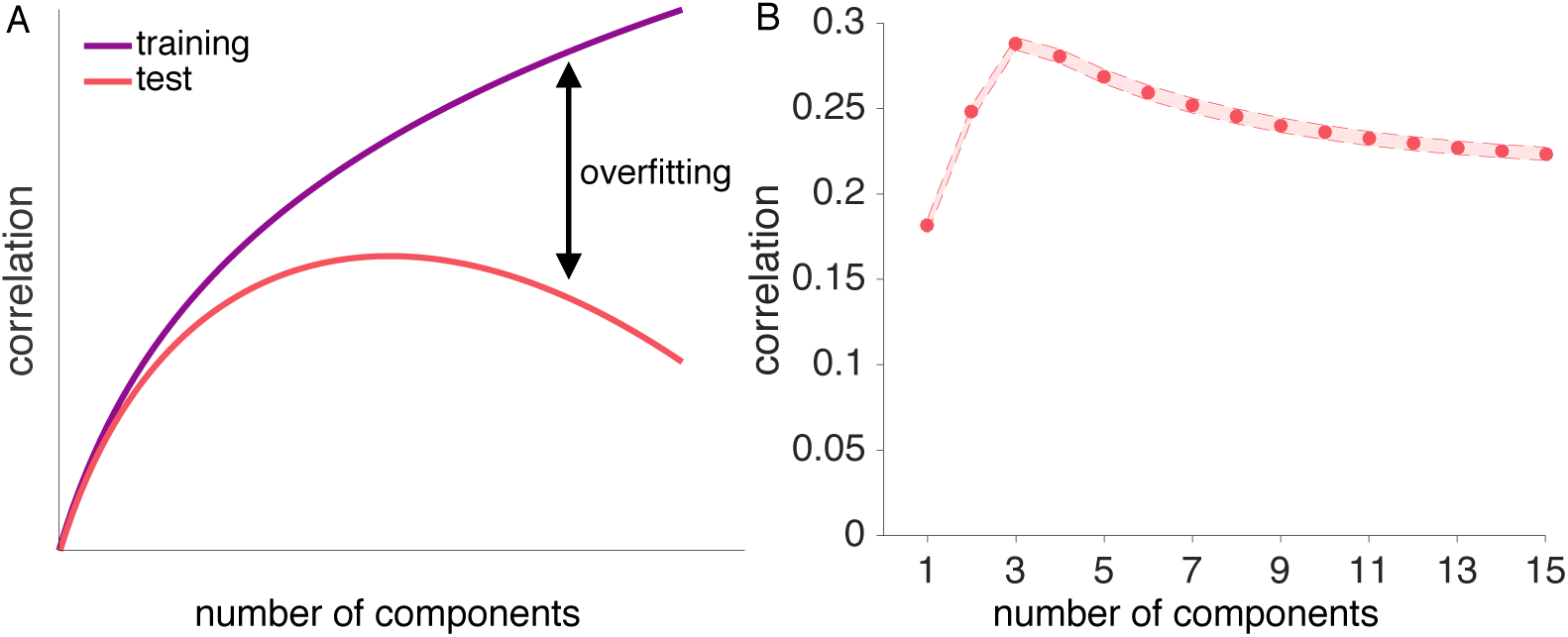
Illustration of the concept of overfitting and generalizability. A: As more components are added to a low-dimensional reconstruction, the correlation between the training data and the reconstruction approaches the maximum of 1 for a full-dimensional reconstruction (purple curve). Adding components is equivalent to adding model parameters to improve fit, which reduces the model’s bias and increases its variance. For the correlation between the reconstructed training and independent test data (red curve), adding components initially improves performance but at some point reduces performance due to overfit (see Parpart et al., 2017, for a related illustration). B: Reconstruction correlations achieved by all possible low-dimensional reconstructions for a simulated ground-truth dimensionality of 4. Reconstruction correlations rise as more components are added up to the point where the true dimensionality is reached, and decrease afterwards. Results are averaged across 6 runs and 1000 simulated voxel patterns.

One naive way to navigate this trade-off between simple and complex models is to use some arbitrary cutoff, such as including the number of components that captures some amount of variance in the training data or deciding based on visual inspection which components may carry signal (known as scree plot, Cattell, 1966). In the case of fMRI, where the signal-to-noise ratio depends on multiple factors like scanner settings, experimental design, and physiological activity (Huettel et al., 2003), estimating the underlying dimensionality based on an arbitrary cut-off criterion for explained variance could be misleading. Likewise, although identifying relevant components via visual inspection works for small datasets, it is not applicable to large datasets as fMRI data, as it would require a manual decision for each set of voxels. Furthermore, the size of fMRI datasets (usually thousands of voxels) calls for a computationally efficient and automated approach, making estimating the dimensionality for the whole brain feasible. Thus, for neuroimaging data, there is a need for an efficient, systematic and objective approach that can both identify areas with statistically significant dimensionality and provide a useful estimate of the underlying dimensionality.

Previous efforts to estimate the dimensionality of neural response patterns have applied linear classifiers to neural data to evaluate dimensionality (Rigotti et al., 2013; Diedrichsen et al., 2013). Rigotti et al. (2013) were able to show that dimensionality of single-cell recordings in monkey PFC is linked to successful task-performance, indicating that dimensionality of neural patterns is task-sensitive. In line with this, Diedrichsen et al. (2013) showed that the dimensionality of motor cortex representations differs depending on the task. Using a combination of PCA and linear Gaussian classifiers, the authors showed that motor cortex representations of different force levels are low dimensional, whereas usage of different fingers was associated with multidimensional neural patterns (Diedrichsen et al., 2013). Notably, both studies focused on estimating task-related changes in dimensionality in a prescribed brain region, rather than estimating which areas across the brain had significant dimensionality. Other methods test dimensionality solutions against a noise distribution constructed by permuting the original data Lehky et al. (2014). However, such methods do not respect the spatial and temporal correlation structure in fMRI data as our method does. Although these methods highlight the potential to estimate the dimensionality of a neural pattern in a prescribed region, they are computationally demanding and require close inspection of the results, which can be impractical in situations such as in a searchlight analysis.

In the present work, we expand on previous contributions by evaluating a novel approach that, in a robust and computationally efficient manner, tests which areas display statistically significant dimensionality, estimates the dimensionality, and provides an indication of the uncertainty of the estimate.

We combine singular value decomposition (SVD) and cross-validation to identify areas across the brain with underlying dimensionality. We derive which of all possible low-dimensional reconstructions of the fMRI signal is the best dimensionality estimate of a held-out test run, and quantify the goodness of the low-dimensional reconstruction via Pearson correlation.

Using a cross-validation procedure to identify the best dimensionality esimate boosts that only components that carry signal and thus generalize to new data are kept. By assessing the significance of the correlation, we can distinguish between areas that show reliable signal with underlying dimensionality vs. areas that do not show a reliable task-modulation. We will refer to this task-dependent dimensionality as functional dimensionality. After establishing significant functional dimensionality, we use Bayesian modeling to derive a population estimate and associated uncertainty of the degree of dimensionality.

We define functional dimensionality as reliable task-dependent changes in a neural pattern that generalize across runs within a subject, though the representational geometry need not be common across subjects. A prerequisite for functional dimensionality is that neural patterns are reliable within subjects. As we show below (see also Figure 1B), our approach can find the low-dimensional projection of a neural pattern that generalizes best across runs.

Through simulations and evaluation of three (published) fMRI datasets, we find that our method successfully identifies areas with significant functional dimensionality and provides reasonable estimates of the underlying dimensionality. In the first fMRI dataset, participants performed a categorization task which required differential attention to various stimulus features (Mack et al., 2013). The second study investigated shape- and category specific neural responses to the presentation of natural images (Bracci and Op de Beeck, 2016). The third study involved categorization tasks that varied systematically in their attentional demands (Mack et al., 2016), which we predict should affect functional dimensionality.

Across all three studies, we were able to identify areas carrying functional dimensionality in a manner that supported and extended the original findings. Focusing on wholebrain effects in the the first two studies, we identified a consistent network of areas showing functional dimensionality during visual stimulus processing. This network encompassed areas that were reported by the original authors as being task-relevant, identified through representational similarity analysis and cognitive model fitting (Bracci and Op de Beeck, 2016; Mack et al., 2013). Furthermore, functional dimensionality was revealed in additional areas, highlighting the sensitivity of our method and suggesting that reliable task-modulated signal was present that was not explained by the models the original authors tested. In the last study, we combined a region-of-interest approach and multilevel Bayesian modeling to show that dimensionality varied depending on task-requirements, which follows from the original authors’ claims but remained untested until now (Mack et al., 2016). We outline how the notion and identification of functional dimensionality can aid the analysis and understanding of neuroimaging data in various ways.

## 2. General Methods

Neuroimaging data, such as fMRI, M/EEG, or single-cell recordings, can be represented as a matrix of *n* voxels, neurons, or sensors *× m* conditions. For example, BOLD response patterns in the fusiform face area (FFA) to 3 different stimulus conditions can be expressed as a matrix *Y* of the size *n* (number of voxels) × 3 (face, house, or tool stimulus condition). The maximum possible dimensionality is determined by the minimum of *n* and *m*, which in this example would be 3, assuming many voxels in FFA were included in the analysis. As fully explained below, the maximum possible dimensionality is *m* – 1 (in this example, 3 – 1 = 2) because each voxel (i.e., matrix row) is mean-centered. In this toy example, rest is implicitly included as a condition, that is, even if all conditions showed the same activity pattern, the estimated dimensionality would be 1. Mean-centering the voxel patterns beforehand accounts for this.

However, functional dimensionality could be lower. For example, dimensionality would be lower if the region only responded to face stimuli and showed the same lower response to house and tool stimuli.

The approach to dimensional estimation we present here is modular and estimates a matrix’s dimensionality by combining low-rank approximation with cross-validation and significance testing. This modularity allows to flexibly choose the dimensionality reduction technique which best fits with ones requirements. Here, we used SVD (which is often used to compute PCA solutions) because it is a well-understood, easy to implement, and a computationally efficient low-rank matrix approximation.

The choice of SVD, as well as how the data matrix is normalized is informed by our understanding of the underlying neural signal. Because voxels differ greatly from one another in their overall activity level and activity levels can drift over runs, we mean-center each row (i.e., voxel) of the data matrix by run. In contrast, we do not mean-center each column, as would typically be done with approaches that focus on the covariance of the column vectors (e.g., PCA). The reason we do not normalize by column (i.e., condition) is that we are open to the possibility that different stimuli may be partially coded by overall activity levels of a population of voxels. For example, imagine a brain area only responds strongly to faces, but not to other stimuli. An SVD with demeaned voxels (i.e., rows) would be sensitive to this dimension of representation, whereas a procedure that effectively worked with demeaned columns would not be sensitive to this task-driven difference in neural activity (see Davis et al., 2014; Hebart and Baker, 2017; Diedrichsen and Kriegeskorte, 2017, for a related discussion).

In the following section, we describe how a combination of SVD and cross-validation can be used to test whether an observed neural pattern can be successfully reconstructed using a low-rank approximation, assessed as a significant Pearson correlation between a low-rank approximation and a held out test set, and how this technique provides an estimate of the pattern’s underlying dimensionality (see Figure 2 for an overview of all steps). As all our examples are fMRI data sets, we will describe the steps using fMRI terminology, though the procedure could be applied to any type of neuroimaging data. We provide the code and data to replicate the analyses presented here and for use on other datasets at osf.io/tpq92.

**Figure 2.**
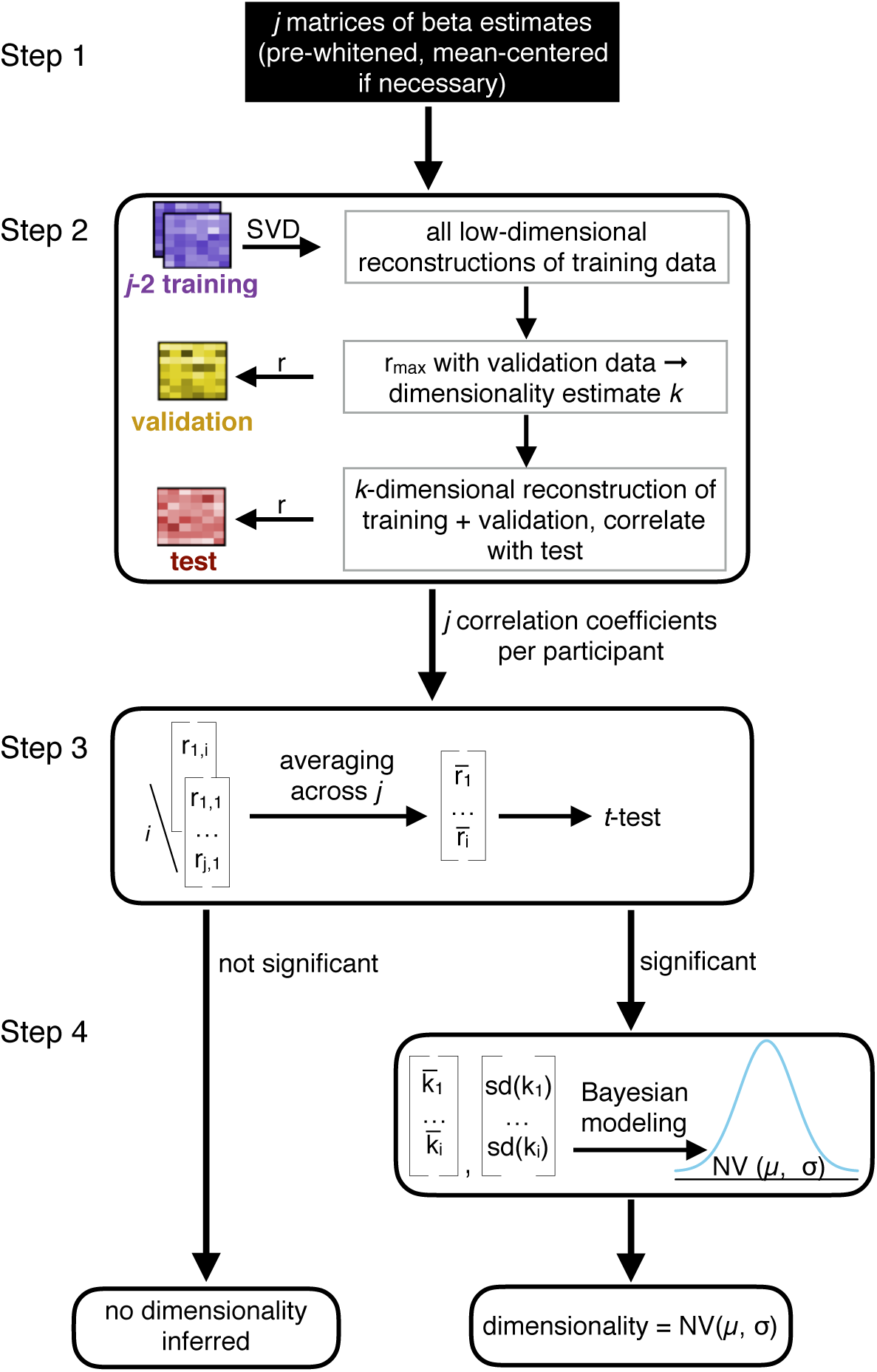
*(previous page)*: Step 1: Prior to dimensionality estimation, raw data are pre-processed with preferred settings and software and beta estimates derived from a GLM are obtained for each condition of interest. The resulting *j* matrices of size *n* (number of voxels) *×m* (number of conditions) are pre-whitened and mean-centered (by row, i.e., voxel) to remove baseline differences across runs. Step 2: a combination of cross-validation and SVD is implemented to find the best dimensionality estimate *k* for each run *j*. Pearson correlations between all possible low-dimensionality reconstructions of the data and a held-out test set quantify the goodness of each reconstruction for each run *j* (see Figure 3 for details). Step 3: the resulting *j* correlations are averaged for each participant and tested for significance, for instance using *t*-tests, across all participants. Step 4: If the reconstruction correlations are significant across participants, a hierarchical Bayesian model can be used to derive the best estimate of the degree of functional dimensionality (see Figure 4 for details). For each participant, the average estimated dimensionality and standard deviation of this estimate is calculated and a population estimate and respective standard deviation (uncertainty in the estimate) is derived across all participants.

### 2.1 Step 1: Data pre-processing

We developed the presented method with application to fMRI data in mind, though it can be easily adapted to fit requirements of single cell recordings or M/EEG data. The method takes beta estimates resulting from a GLM fit to the observed BOLD response as input. In all studies presented here, standard pre-processing steps were performed using SPM 12 (Wellcome Department of Cognitive Neurology, London, United Kingdom), but the precise nature of the preprocessing and implemented GLM is not critical to our method. Functional data were motion corrected, co-registered and spatially normalized to the Montreal Neurological Institute (MNI) space.

To reduce the impact of the structured noise, which is correlated across voxels, on the dimensionality estimation and to improve the reliability of multivariate voxel response patterns (Walther et al., 2016), we applied multivariate noise-normalization, that is, spatial pre-whitening, before estimating the functional dimensionality. We used the residual time-series from the fitted GLM to estimate the noise covariance Σ_*noise*_ and used regularization to shrink it towards the diagonal (Ledoit and Wolf, 2004). Each *n × m* matrix of beta estimates *Y* was then multiplied by 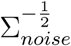 (Walther et al., 2016).

In fMRI data, the baseline activation can differ across functional runs. This has important implications for our approach presented here, as it can bias the correlation between neural patterns across runs. To account for this, we demeaned the pre-whitened beta estimates across conditions, resulting in an average estimate of zero for each voxel. This demeaning reduces the possible maximum dimensionality of the data to *k*_*max*_ = *m* − 1. Notably, demeaning of voxels is conceptually different from demeaning conditions, which would have been implemented by PCA, as it preserves differences between conditions, whereas PCA would remove those.

### 2.3 Step 2: Evaluating all possible SVD (dimensional) models

The dimensionality of a matrix is defined as its number of non-zero singular values, identified via singular value decomposition (SVD). SVD is the factorization of an observed *n × m* matrix *M* of the form *U* Σ*V*^T^. *U* and *V* are matrices of size *m × m* and *n × n*, respectively, and Σ is an *n × m* matrix, whose diagonal entries are referred to as the singular values of *M*. A *k*-dimensional reconstruction of the matrix *M* can be achieved by only keeping the *k* largest singular values in Σ and replacing all others with zero, resulting in 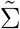. This is known as Eckart-Yong theorem (Eckart and Young, 1936), leading to equation 1:

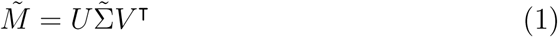

To estimate the dimensionality of fMRI data, we applied SVD to *j*(number of runs) matrices *Y* of *n*(number of voxel) *× m*(number of beta estimates), with the restriction of *n > m*.

Critically, fMRI beta estimates are noisy estimates of the true signal. In the presence of noise, all singular values of a matrix will be non-zero, requiring the definition of a cut-off criterion to assess the number of singular values reflecting signal. Removing noise-carrying components from a matrix is beneficial, as it avoids overfitting to the noise and thus, improves the generalizability of the low-dimensional reconstruction to another sample (see Figure 1 A for an illustration of the concept of overfitting). We aimed to avoid any subjective (arbitrary) criterion as percentage of explained variance or alike (Cattell, 1966). To that end, we implemented a nested cross-validation procedure at the core of our method to identify singular values that carry signal (see step 1 of the general overview depicted in Figure 2 and Figure 3 for a detailed illustration of the cross-validation approach). This allows us to reduce the inflation of dimensionality of fMRI data due to noise and test which areas of the brain carry signal with functional dimensionality.

**Figure 3:**
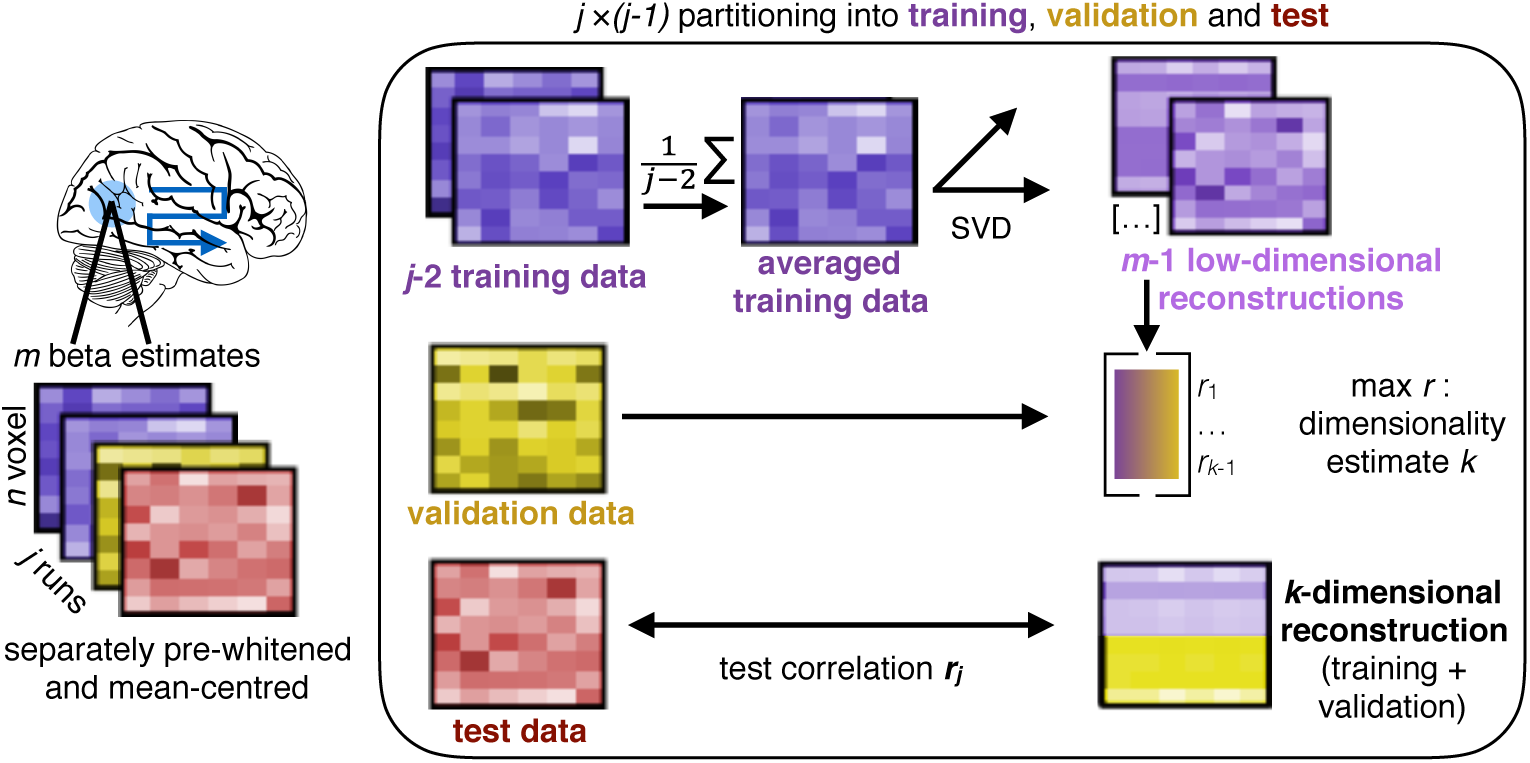
Illustration of the combination of SVD and cross-validation, corresponding to step 2 in Figure 2. For each searchlight or ROI, *j* (number of runs) *n* (number of voxels) *× m* (number of beta estimates) matrices are used to estimate the functional dimensionality. For all possible partitions of *j* runs into training, validation and test data, we first average all training runs and build all possible low-dimensional reconstructions of these averaged data using SVD. All reconstructions are then correlated with the validation run, resulting in *j −* 1 correlation coefficients and respective dimensionalities. The dimensionality that results in the highest average correlation across *j −* 1 runs is picked as dimensionality estimate *k* for this fold and a *k*-dimensional reconstruction of the average of the training and validation runs is correlated with a held-out test-run, resulting in a final reconstruction correlation. In total, *j* reconstruction correlations are returned that can be averaged and tested for significance across participants using one-sample t-tests or alike. To derive a better estimate of the underlying dimensionality, the *j* dimensionality estimates per participant can be submitted to the hierarchical Bayesian model (step 4 in Figure 2)

Data are partitioned *j ×* (*j −* 1) times into training (*Y*_*train*_), validation (*Y*_*val*_), and test (*Y*_*test*_) data. The (demeaned and pre-whitened) *j* − 2 training runs are averaged, and SVD is applied to the resulting *n × m* matrix 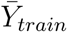. We then build all possible low-dimensional reconstructions of the averaged training data, with dimensionality ranging from 1 to *m* − 1. Low-dimensional reconstructions are generated by keeping only the *k* highest singular values and setting all others to zero. Each low-dimensional reconstruction of matrix 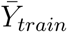 is correlated with the held-out *Y*_*val*_. This is repeated for each possible partitioning in training and validation, resulting in *j* − 1 × *m* − 1 correlation coefficients. Correlations are Fisher’s z-transformed and averaged across the *j* − 1 partitionings. The dimensionality with the average highest correlation is picked as best estimate *k* of the underlying dimensionality. As keeping components that reflect noise rather than signal lowers the correlation with an independent data set, the highest correlation is not necessarily achieved by keeping more components. This procedure thus avoids inflated dimensionality estimates.

After identifying the best dimensionality estimate *k* for run *j*, the training and validation runs from 1 to *j* − 1 are averaged together and SVD is applied to the averaged data. We then generate a *k*-dimensional reconstruction of the averaged data. The quality of this final low-dimensional reconstruction is measured as Pearson correlation with *Y*_*test*_. We chose Pearson correlation instead of mean-square error (MSE) because Pearson correlation is scale invariant.

### 2.3 Step 3: Determining statistical significance

The approach results in *j* estimates of the underlying dimensionality and *j* corresponding test correlations per participant. Under the null-hypothesis of no dimensionality, and thus, only noise present in the matrix, reconstruction correlations averaged across runs are distributed around zero. Thus, across-participants significance of the averaged reconstruction correlations can be assessed using one-sample *t*-tests or non-parametric alternatives, as for instance permutation tests (Nichols and Holmes, 2003), and established correction methods for multiple comparisons, like threshold-free cluster enhancement (TFCE, see Smith and Nichols, 2009).

Only when a significant, *k*-dimensional, reconstruction correlation is found across participants, do we refer to an area as showing functional dimensionality. It should be noted that a significant reconstruction correlation only indicates that the underlying functional dimensionality is one or bigger.

More evidence for a dimensionality of two or larger can be gathered by removing not only the voxel-mean before estimating the dimensionality, but also the condition mean, which removes a potential source of univariate differences between conditions. However, as discussed in Davis et al. (2014) and Hebart and Baker (2017), this does not indubitably mean that the dimensionality of the pattern is two or larger.

### 2.4 Step 4: Estimating the degree of functional dimensionality

The previously described steps allow us to identify which areas carry reliable signal with functional dimensionality, but do not provide a precise estimate of the degree of the underlying dimensionality. The best population estimate of a region’s functional dimensionality should optimally combine information across participants, giving more weight to participants with more reliable estimates, and should furthermore reflect how peaked the distribution of underlying population estimates is, accounting for the fact that different participants could express different true dimensionality.

Given a significant reconstruction correlation across participant, *j* estimates of the degree of dimensionality are obtained (for each voxel, i.e. center of a searchlight, or ROI) for each participant. In a noise-free scenario, all *j* estimates reflect the true dimensionality and thus, direct inference could be made solely based on these estimates. Under noise, these estimates could over- or underestimate the true dimensionality. The less reliable the *j* dimensionality estimates, the higher the variance across them. Mere averaging of the *j* estimates across participants would discard this information, weighting all participants equally, irrespective of their reliability. Down-weighting the influence of less reliable dimensionality estimates on the population estimate leads to a better population estimate (Kruschke, 2014).

To account for this, we implemented a multilevel Bayesian model using the software package Stan (The Stan Development Team, 2017). Given the mean and standard deviation of *j* dimensionality estimates per participant, the model derives the best estimate for the true degree of dimensionality across all participants. Due to the nature of the multilevel model, individual estimates are subject to shrinkage towards the estimated population mean, and the degree of shrinkage is more pronounced for estimates with higher variance and stronger deviation from the estimated population mean (Kruschke, 2014).

Additionally to the estimate of the population dimensionality, the model returns estimates for the population dimensionality’s variance, reflecting the uncertainty of the dimensionality estimate.

For each individual participant, the model estimates the participant’s true underlying dimensionality and returns the uncertainty of this estimate. Though not our focus here, individual differences in dimensionality estimates could be linked to other measures, such as task performance.

#### 2.4.1. Model parametrization

As can be seen in Figure 4, the Bayesian model has four levels: the prior distributions, the population distributions, the individual distributions, and the observed estimates. Apart from the bottom level, that is, the observed estimates, distributional assumptions must be made. For each individual participant, *j* dimensionality estimates are observed. Those reflect noisy estimates of a participant’s true underlying dimensionality. We chose a truncated *t*-distribution as parametrization of the level of the true individual dimensionalities. The parameters of this distribution were the participant’s estimate of the true underlying dimensionality *µ*_*i*_, subject to shrinkage due to other participants’ estimates and the individual’s standard deviation of dimensionality estimates. The truncated *t*-distribution can account for the limited range of possible data points, as there is a natural maximum and minimum dimensionality that could be observed. It furthermore reflects the assumption that under noise, the true dimensionality of an observed pattern, that is, the mean of the *t*-distribution, would still have the highest probability of estimation, with the dispersion of the distribution depending on the number of observations made (here, runs). On the population level, we chose a truncated normal distribution with mean of *µ* and a standard deviation of *σ*, limited to the range of the possible dimensionality estimates. This was chosen to reflect the assumption that participants from the same population should have similar, though not necessarily identical functional dimensionalities. The combination of a normal distribution on the population level and a truncated *t*-distribution on the single subject level ensured that participants with largely dividing dimensionality estimates are shrunk towards the mean of the overall sample in an optimal way.

**Figure 4:**
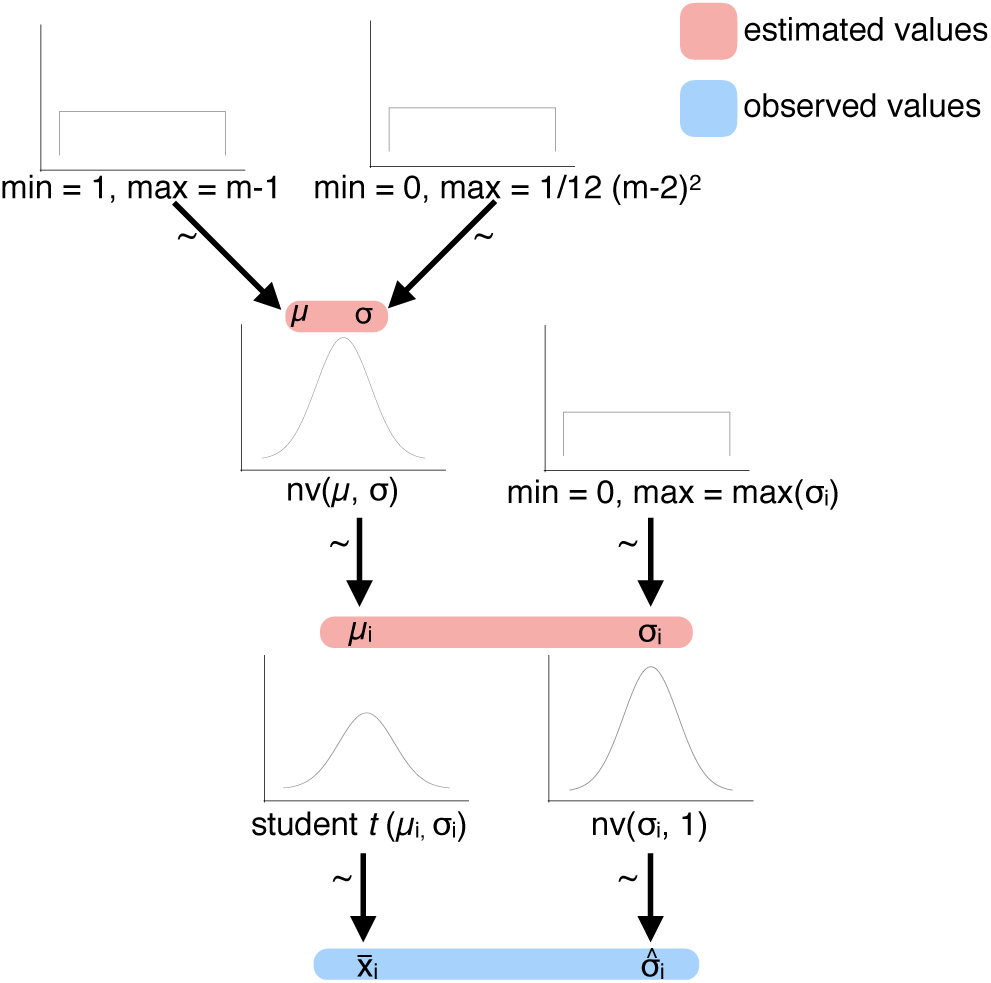
Illustration of the implemented multilevel model to estimate the degree of functional dimensionality, corresponding to step 4 in Figure 2. The observed averaged dimensionality estimates per participant are assumed to be sampled from an underlying subject-specific t-distribution with mean µ_*i*_ and standard deviation σ_*i*_. The standard deviation 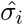 of the participants’ dimensionality estimates is assumed to be sampled from a normal distribution with mean σ_*i*_ and a standard deviation of 1. The subject-specific *t*-distributions of µ_*i*_ are assumed to come from a population distribution with a normally distributed mean *µ* and variance *σ*. Subject-specific standard deviations σ_*i*_ are assumed to come from a uniform distribution, ranging from 0 to max(σ_*i*_). At the top level, a uniform prior is implemented. Mean and variance of the normal distribution of population means *µ* are assumed to come from a uniform distribution ranging from 1 to *m −* 1 and 0 to σ_*max*_, respectively. Distributions were derived from https://github.com/rasmusab/distribution_diagrams.

As we did not have strong priors regarding the dimensionality of the neural patterns, we implemented a uniform prior over the population dimensionality estimates, reflecting that the dimensionality could be anything from 1 to *m* − 1.

Notably, this does not imply that all participants need to show an estimated functional dimensionality larger than zero, but rather reflects the assumption that a significant second-level functional dimensionality suggests a non-zero functional dimensionality in the population.

The prior distribution can be adapted to be informative for studies estimating the functional dimensionality of neural patterns with stronger priors. Figure 4 shows an illustration of the model.

The model is formally expressed in Equation 2.

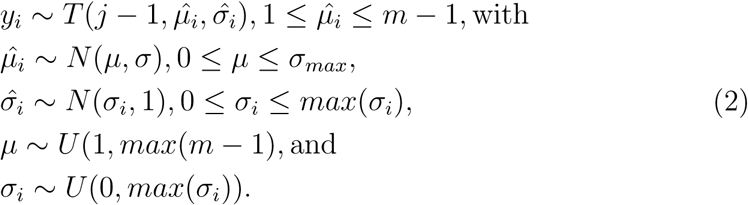

The maximum population variance was defined as the expected variance of this uniform distribution 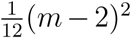, reflecting the prior that each participant could express a different, true dimensionality. On the subject-level, the maximum variance was defined as

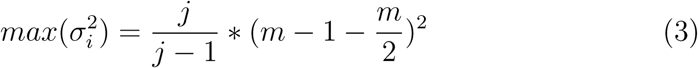

which corresponds to the maximum possible variance across *j* dimensionality estimates.

The *j* estimates of a participant’s dimensionality were not independent, since the training data overlapped. Thus, the standard deviation of the estimates will be underestimated. The degree of this underestimation will be the same for all participants though, which allows us to rely on the observed standard deviation as a proxy for the estimation noise without correcting.

## 3. Simulations

Before applying our method to real fMRI data, we tested the validity of our method through dimensionality-recovery studies on simulated fMRI data. Estimating the dimensionality for simulated cases where the true underlying dimensionality is known allowed us to assess whether our procedure results in a reliable dimensionality estimate.

### 3.1 Methods

Simulated data were created using the RSA toolbox (Nili et al., 2014) and custom Matlab code. Parameters of the simulation were picked in accordance with the study by Mack and colleagues (2013). We simulated fMRI data of presentation of 16 different stimuli, presented for 3 sec, three repetitions per run, and six runs, closely matching the specifications of the original study. To mimic a searchlight-approach, we defined the size of the cubic sphere 4 × 4 × 4 voxels, resulting in a simulated pattern of 64 voxels.

We simulated data with a dimensionality of 4, 8, and 12 and ten steps of exponentially increasing noise levels to investigate how noise affects dimensionality estimates, and how this effect interacts with the ground-truth dimensionality. In order to apply hierarchical Bayesian model, we created simulated data for 20 ‘participants’. For each simulated participant, the noise level was drawn from a normal distribution (truncated at 0.5 and 2 times the average noise level).

To generate data with varying ground-truth dimensionality *k*, we first generated true, i.e. noise-free, *n*(voxel) × *m*(conditions) matrices with underlying pre-defined dimensionality. This was achieved by applying PCA to a random 16 × 16 matrix and building a *k*-dimensional reconstruction of it. All eigenvalues of this initial *k*-dimensional matrix had the same value. Rows of this matrix were added to an *n* × 16 matrix. For each row, i.e. voxel, a specific amplitude was drawn from a normal distribution and added.

In the next step, we calculated the dot-product of the generated beta matrices and generated design matrices, which were HRF convolved. This resulted in noise-free fMRI time series.

A noise matrix was generated by randomly sampling from a Gaussian distribution. The *n*(voxel) × *t*(timesteps) matrix was then spatially smoothed and temporally smoothed with a Gaussian kernel of 4 FWHM. Finally, this temporally and spatially smoothed noise matrix was added to the noise-free time-series and the design matrix was fit to the resulting data using a GLM. This resulted in a (noisy) voxel × conditions beta matrix for each simulated run. The generated beta matrices were then passed on to the dimensionality estimation.

To gather a reliable estimate of the performance of our procedure, we ran a total of 100 of these simulations for each combination of ground-truth dimensionality and noise-level.

We then estimated the dimensionality for each simulated participant as described above and passed each participant’s average estimated dimensionality and the standard deviation of this estimate to the described hierarchical Bayesian model. To assess the goodness of the estimated dimensionalities, we combined all posterior estimates of the single simulated participants’ dimensionalities (parameter µ_*i*_) across all simulated voxels. The width of the distributions of these posteriors reflects the uncertainty of the estimated population dimensionality, and the distributions’ means reflect the estimated population dimensionality.

### 3.2 Results and Discussion

Across 100 simulations of data with a ground-truth dimensionality of 4, 8, or 12 and ten different noise levels, we assessed how estimated dimensionalities are affected by noise and how this effect interacts with the ground-truth dimensionality.

Ideally, our method would exhibit these properties: 1) The posterior estimate of the degree of underlying dimensionality should be close to the ground-truth dimensionality when the signal-to-noise ratio is high. 2) The uncertainty of the posterior estimate should increase with increasing noise-levels. 3) Estimates should gracefully degrade such that as noise increases the relative order of ground-truth dimensionalities should still be reflected in the estimated dimensionalities and the posteriors still contain the ground-truth values. 4) With increasing noise, the relative importance of the prior should increase and in the limit all ground-truth dimensionalities should converge to the mean of the prior.

As can be seen in Figure 5A, the results from the simulation show that our method meets all four criteria. For a low noise level, the estimated dimensionalities largely overlap with the ground-truth and are very consistent across simulated participants. With increasing noise, estimated dimensionalities deviate more strongly from the underlying ground-truth and move towards the mean of the uniform prior. Furthermore, the uncertainty in the dimensionality estimates increases, reflected in the width of the distributions.

**Figure 5:**
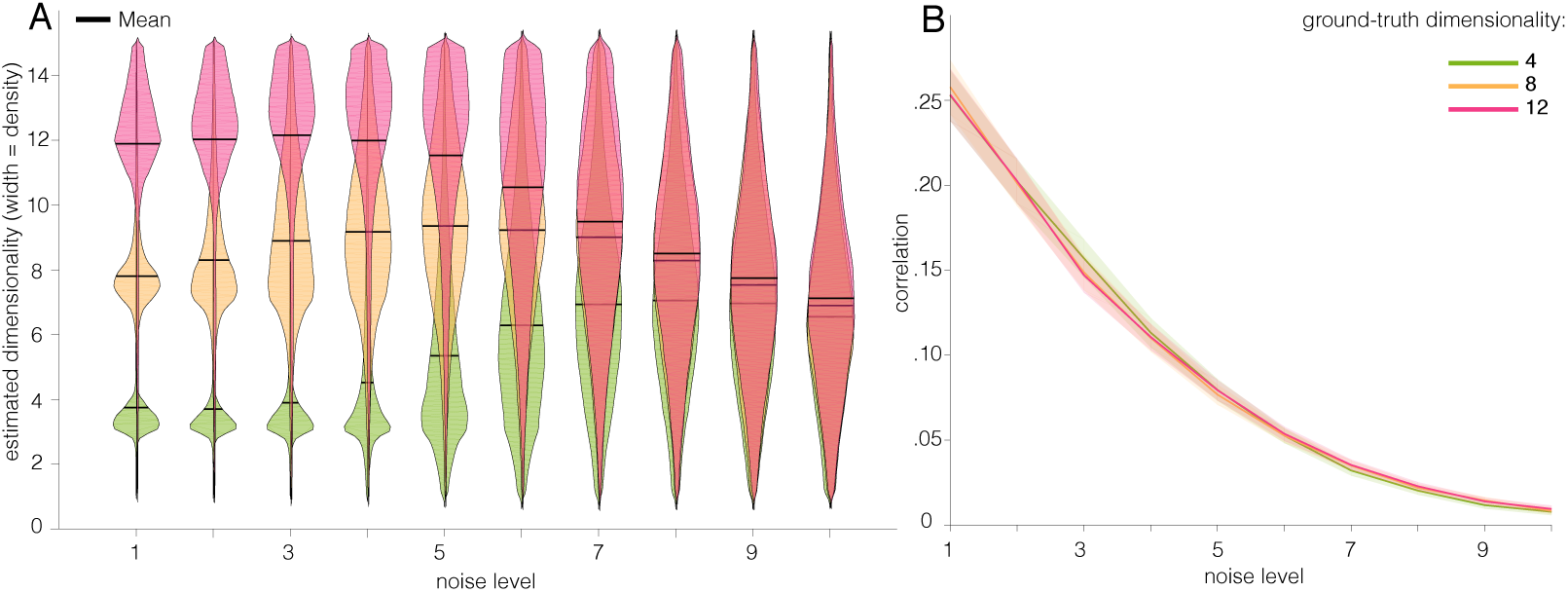
Results from the simulation. A: Distributions of single-subject posterior dimensionality estimates for a ground-truth dimensionality of 4, 8, or 12 and increasing noise levels. As noise increases, the estimates become less accurate and less certain, as indicated by the width of the distributions. For the highest noise level, the posterior distributions for all ground-truth dimensionalities overlap largely. B: Average reconstruction correlations for the different ground-truth dimensionalities and increasing noise levels. As the noise level increases, reconstruction correlations drop, and this effect is the same across the three different ground-truth dimensionalities.

Figure 5B shows the average reconstruction correlations with the held out test data for the different ground-truth dimensionalities and the different noise levels, which are highly overlapping.

An additional observation from the simulation results is that moderate levels of noise can lead to a small inflation of estimated dimensionalities for higher ground-truth dimensionality levels, as seen here for the case of a dimensionality of 8. This effect is due to the correlational structure of noise in fMRI data. The SVD is sensitive to this correlational structure and as a result, singular values that reflect noise could surpass singular values that reflect signal. This would then cause an overestimation of the underlying dimensionality, since keeping a noise-carrying singular component would not improve the correlation with the held-out validation data, but adding the next, signal-carrying singular value to the reconstruction would. However, this inflation is only minor. It does not violate the rank order of the posterior dimensionality estimates and the distribution of the posterior dimensionality estimates reflects the increased uncertainty in the estimate.

Together, these simulations show that our procedure is suitable to provide an accurate estimate of the degree of underlying functional dimensionality for good signal-to-noise ratios. Moreover, the access to the whole distribution of dimensionality estimates allows to draw valid inferences on the relative degree of functional dimensionality even under high noise, and the width of the distribution of these estimates reflects the uncertainty of these estimates. Thus, the combination of cross-validated SVD and hierarchical Bayesian modeling can provide a robust and interpretable estimated distribution of the degree of underlying functional dimensionality, which reflects the certainty in the estimate.

## 4. Data sets

Following the successful tests of our procedure with simulated data, we applied our method to three different, previously published fMRI datasets, all employing visual stimuli and testing healthy populations. We tested three core aims of our method: 1) Identifying areas carrying functional dimensionality, 2) Using functional dimensionality to assess sensitivity to stimulus features, and 3) Measuring task-dependent differences in dimensionality.

### 4.1 Identifying areas carrying functional dimensionality

Using data from a category learning study by Mack et al. (2013), we aimed to identify areas carrying functional dimensionality and compare them with the areas found by the original authors’ model-based analysis. Model-based analyses test specific assumptions about representational geometry that our approach does not. Furthermore, these analyses require some underlying dimensionality to identify an area. Therefore, we expected our method to reveal significant functional dimensionality in all areas that were reported in the original study, as well as additional areas that were reliably modulated by the task in a way that was not captured by the model tested in the original publication.

#### 4.1.1. Methods

Participants were trained on categorizing nine objects that differed on four binary dimensions: shape (circle/triangle), color (red/green), size (large/small), and position (left/right). During the fMRI session, participants were presented with the set of all 16 possible stimuli and had to perform the same categorization task. Out of 23 participants, 20 were included in the final analysis presented here, with 19 participants completing 6 runs composed of 48 trials and one participant completing 5 runs.

Standard pre-processing steps were carried out using SPM12 (Penny et al., 2006) and beta estimates were derived from a GLM containing one regressor per stimulus (16 in total, see Supplemental Materials for details). The dataset was retrieved from osf.io/62rgs.

We ran a whole-brain searchlight with a 7mm radius sphere and a voxel size of 3 × 3 × 3mm to estimate which brain areas carry signal with functional dimensionality, that is, signal that could be reliably predicted across runs based on a low-dimensional reconstruction. For each searchlight, data were pre-whitened and mean-centered as described above. Dimensionality estimation was performed as previously described and the resulting *j* correlations and dimensionality estimates were ascribed to the center of the searchlight. The code for the searchlight was based on the RSA toolbox (Nili et al., 2014).

For each voxel, the *j* correlation coefficients were averaged and their significance was assessed via non-parametric one-sample t-tests across subjects using FSL’s randomise function (Winkler et al., 2014). Results were family-wise error (FWE) corrected using a TFCE threshold of *p* < .05.

In their original analysis, the authors fit a cognitive model to participants classification behavior to estimate attention-weights to the single stimulus features. Based on these attention weights, they derived model-based similarities between stimuli and used RSA to examine which brain regions show a representational geometry that matches with these predictions. We replicated this analysis using the same beta estimates that were passed on to the dimensionality estimation in order to maximize comparability of the two approaches. As for estimating the dimensionality, we ran a whole-brain search-light with a 7mm radius sphere (based on the RSA toolbox, Nili et al., 2014). We averaged voxel response patterns across runs and calculated the representational distance matrices (RDM) as all pairwise 1–Pearson correlation distance. We assessed correspondence of these RDMs with the model-based distance matrices via Spearman correlation. The resulting Spearman correlation for each participant was assigned to the center of the searchlight and their significance was assessed via non-parametric one-sample t-tests across subjects using FSL’s randomise function (Winkler et al., 2014). Results were family-wise error (FWE) corrected using a TFCE threshold of *p* < .05.

#### 4.1.2. Results

We aimed to identify areas that show functional dimensionality and examine how those overlap with the authors’ original findings implementing a model-based analysis. We found significant dimensionality (i.e., reconstruction correlations) in an extended network of occipital, parietal and prefrontal areas (see Figure 6). In these areas, signal was reliable across runs and showed functional dimensionality.

**Figure 6:**
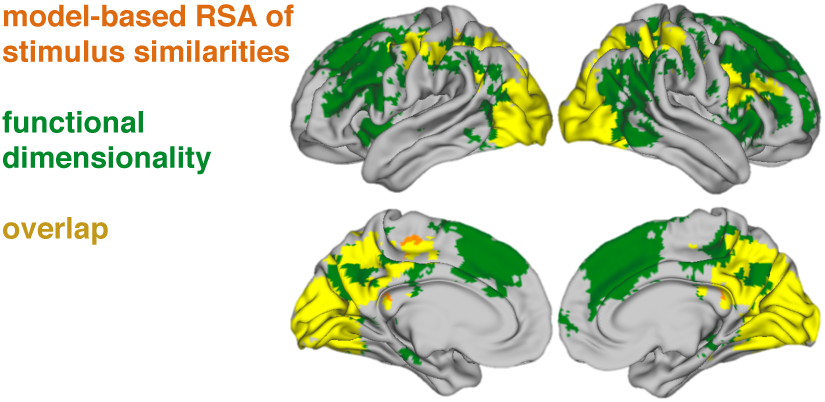
Areas that showed significant functional dimensionality (green), significant fit with the RSA comparing neural representational similarity with model-based predictions of stimulus similarity (orange), or both (yellow). FWE-corrected using a TFCE threshold of *p* < .05. Notably, our method identifies large clusters of functional dimensionality in prefrontal cortex, indicating that areas here were consistently engaged by the task, though their patterns did not fit with the implemented cognitive model.

As can be seen in Figure 6, our method successfully identified all areas that were found in the original model-based analysis, which bolsters the authors original interpretation of their results. Notably, we were able to identify further areas that did not show a fit with the implemented attention-based model, suggesting that signal changes in those areas reflect a different aspect of the task space than captured by the cognitive model.

#### 4.1.3. Discussion

Within the first dataset, we showed that by identifying areas with significant functional dimensionality, it is possible to reveal areas that can plausibly be tested for correspondence with a hypothesized representational similarity structure, as for instance derived from a cognitive model. More specifically, we were able to identify all areas that have been reported in the original analysis by Mack et al. (2013) to show a representational similarity as predicted by a cognitive model. Additionally, we found further areas that had not been revealed in the original analysis to show functional dimensionality. This indicates that those areas have a reliable functional dimensionality but reflect cognitive processes or task-aspects that are not captured by the cognitive model. For instance, activation in the medial BA 8 has been found to correlate with uncertainty and task-difficulty (Volz et al., 2005; Huettel, 2005; Crittenden and Duncan, 2014), suggesting that the neural patterns in this region in the current task might reflect processes related to the difficulty or category uncertainty of the categorization decision for each stimulus. Given that our method identifies more areas than model-based RSA, one might be tempted to view it as a more powerful and statistical sensitive version of RSA, but such an interpretation would be incorrect. Whereas RSA evaluates specific assumptions regarding representational geometry, tests of functional dimensionality depend solely on reliability of patterns (assessed across runs). Together, the findings highlight the potential of our procedure to aid evaluation of model performance and identify areas ahead of model-fitting.

### 4.2 Using functional dimensionality to assess sensitivity to stimulus features

Using data from a study with real-world categories and photographic stimuli by Bracci and Op de Beeck (2016), we tested whether different brain regions show functional dimensionality in response to different stimulus groupings (i.e., depending on how the stimulus-space is summarized). For example, the columns in the data matrix may be organized along either visual categories or shape. In this fashion, our technique could be useful in evaluating general hypotheses regarding the nature and basis of the functional dimensionality in brain regions.

#### 4.2.1. Methods

During the experiment, participants were presented repeatedly with 54 different natural images that were of nine different shapes and belonged to six different categories (minerals, animals, fruit/vegetables, music instruments, sport instruments, tools), allowing the authors to dissociate between neural responses reflecting shape or category information.

Standard pre-processing of the data was carried out using SPM12 (see Supplemental Material for details). In line with the authors original analysis, we tested for differences depending on whether the stimuli were averaged to emphasize their category or shape information. To that end, we constructed two separate GLMs. The first GLM (catGLM) was composed of one regressor per category (six in total), thus averaging across objects shapes. The second GLM (shapeGLM) consisted of nine different regressors, one for each shape, averaging neural responses across object categories. In both GLMs, regressors were convolved with the HRF and six motion-regressors as covariates of no interest were included.

Dimensionality was estimated separately for both GLMs. We ran a whole-brain searchlight with a 7mm sphere (voxel size of 3 × 3 × 3*mm*) on the beta estimates of the respective GLM, again pre-whitening and mean-centering voxel patterns within each searchlight before estimating the dimensionality. Reconstruction correlations were averaged across runs for each participant and tested for significance across participants using FSL’s randomise function (Winkler et al., 2014). Results were FWE corrected using a TFCE threshold of *p* < .05.

#### 4.2.2. Results

When testing for functional dimensionality for the shape-sensitive GLM, we found significant reconstruction correlations in bilateral posterior occipitotemporal and parietal regions, indicating functional dimensionality in these areas. Additionally, a significant cluster was revealed in the left lateral prefrontal cortex (see Figure 7). Testing for functional dimensionality for the category-sensitive GLM also revealed strong significant correlations in occipital and posterior-temporal regions, but notably showed more pronounced correlations in bilateral lateral and medial prefrontal areas as well. This is in line with the authors original findings that showed that neural patterns in parietal and prefrontal ROIs correlated more strongly with a model reflecting category similarities, whereas shape similarities were largely restricted to occipital and posterior temporal ROIs.

**Figure 7:**
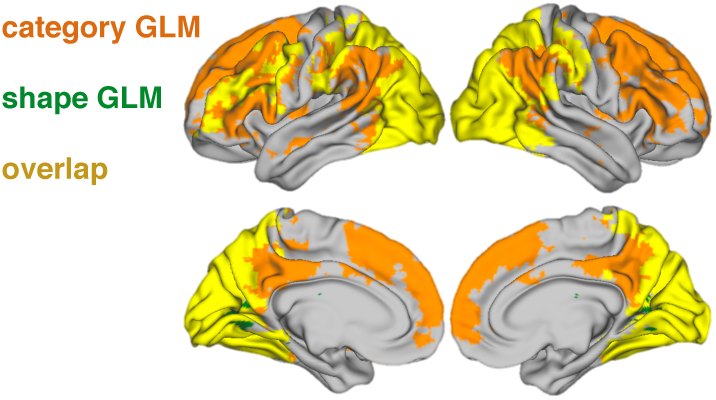
Areas showing significant functional dimensionality for the shape GLM (green), the category GLM (orange), or both (yellow). Results are FWE-corrected using an TFCE threshold of *p* < .05. Across both GLMs, posterior and parietal regions show functional dimensionality. Prefrontal regions show more pronounced functional dimensionality for the category GLM, in line with the original findings.

#### 4.2.3. Discussion

With the second dataset, we tested whether different areas are identified to express significant functional dimensionality depending on how the underlying task-space is summarized. In line with the original authors’ findings (Bracci and Op de Beeck, 2016), we found more pronounced functional dimensionality in prefrontal regions for the GLM emphasizing the category-information across stimuli, compared to the one focusing on shape-information. Likewise, functional dimensionality in occipital regions was more pronounced for the shape-based GLM.

However, compared to the authors’ original findings, we did not find a sharp dissociation between shape and category. For example, we find both shape and category dimensionality present in early visual regions and shape dimensionality extending into frontal areas.

As discussed in the previous section, our method provides a general test of dimensionality whereas the original authors evaluate specific representational accounts that make additional assumptions about shape and category similarity structure. Comparing results suggest that to some degree the dissociation found in Bracci and Op de Beeck (2016) rests on these specific assumptions. A more general test of functional dimensionality, for stimuli organized along shape or category, provides additional information to assist in interpreting the cognitive function of these brain regions, which complements testing more specific representational accounts.

Additional information could be gleamed by estimating differences in dimensionality. In the case of the shape and category GLMs considered in this section, interpretation would be somewhat complicated by the different properties of these two GLMs, including differences in the maximum possible number of dimensions. In the next section, we consider a more straightforward case in which the same GLM is used to compare task influences on functional dimensionality.

### 4.3 Measuring task-dependent differences in dimensionality

In this third dataset, we consider whether the underlying dimensionality of neural representations changes as a function of task. In Mack et al. (2016), participants learned a categorization rule over a common stimulus set that either depended on one or two stimulus dimensions. We predicted that the estimated functional dimensionality, as measured by our hierarchical Bayesian method, should be higher for the more complex categorization problem, extending the original authors’ findings.

#### 4.3.1. Methods

Participants learned to classify bug stimuli that varied on three binary dimensions (mouth, antenna, legs) into two contrasting categories based on trial-and-error learning. Over the course of the experiment, participants completed two learning problems (in counterbalanced order). Correct classification in type I problem required attending to only one of the bugs features, whereas classification in type II problem required combining information of two features in an exclusive- or manner.

Previous research has shown that neural dimensionality appropriate for the problem at hand is linked to successful task performance (Rigotti et al., 2013). Thus, we hypothesized that dimensionality of the neural response would be higher for type II compared to type I in areas known to process visual features, as for instance lateral occipito-temporal cortex (LOC; see e.g. Eger et al., 2008). We included data from 22 participants in our analysis (one participant was excluded due to artifacts in the fMRI data, please refer to the Supplemental Material for further details on the experiment and data preprocessing). The dataset was retrieved from osf.io/5byhb.

In order to infer the degree of functional dimensionality, we estimated it across ROIs encompassing LOC in the left and right hemisphere separately for the two categorization tasks. Because the relevant stimulus dimensions were learned through trial-and-error learning, we excluded the first functional run (early learning) of each problem and analyzed the remaining three runs for each problem.

Prior to estimating the dimensionality, data were pre-whitened and mean-centered. Dimensionality was estimated across all voxels for each ROI and problem, resulting in 3 (runs) × 2 (ROIs) × 2 (problems) correlation co-efficients and dimensionality estimates. Correlation coefficients were averaged per participant, ROI and problem and tested for significance using one-sample *t*-tests. To derive the best population estimate for the underlying dimensionality for each ROI and problem, we implemented the above described hierarchical Bayesian model. To that end, we calculated mean and standard deviation of each participant’s dimensionality estimate per ROI and problem and used those summary statistics to estimate the degree of underlying dimensionality for each ROI and problem.

#### 4.3.2. Results

Estimating dimensionality across two different ROIs in LOC and two different tasks allowed us to test whether the estimated dimensionality differs across problems with different task-demands. As participants had to pay attention to one stimulus feature in the type I problem and two stimulus features in the the type II problem, we hypothesized that dimensionality of the neural response would be higher for type II compared to type I in an LOC ROI.

Both ROIs showed significant reconstruction correlations across both tasks (lLOC, type I: *t*_21_ = 3.08, *p* = .006; rLOC, type I: *t*_21_ = 2.21, *p* = .038; lLOC, type II: *t*_21_ = 3.03, *p* = .006; rLOC, type II: *t*_21_ = 3.37, *p* = .003). This shows that signal in the LOC showed reliable functional dimensionality across runs for both problem types, which is a prerequisite for estimating the degree of functional dimensionality.

To estimate whether the dimensionality differed across problems, we analyzed the data by implementing a multilevel Bayesian model using Stan (The Stan Development Team, 2017), see Figure 2 for an illustration of the model. As hypothesized, the estimated underlying dimensionality was higher for the type II problem compared to type I (type I: µ_*left*_ = 2.92 (CI 95% : 1.33, 4.33), µ_*right*_ = 2.66 (CI 95% : 1.23, 4.14); type II: µ_*left*_ = 4.74 (CI 95% : 3.20, 6.46), µ_*right*_ = 4.69 (CI 95% : 3.56, 6.06), see Figure 8).

**Figure 8:**
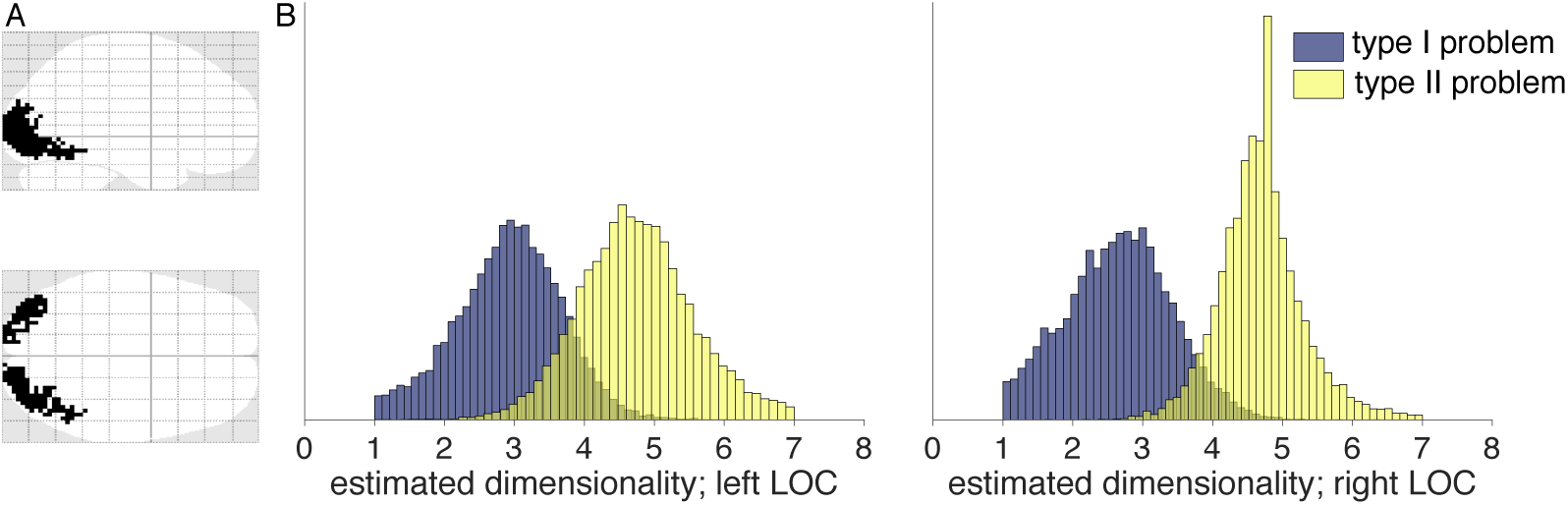
Results of estimating functional dimensionality for two different categorization problems. A: Outline of the two ROIs in left and right LOC. B: Histograms of posterior distributions of estimated dimensionalities in left and right LOC for the type I and II problems. Dimensionalities were estimated by implementing separate multilevel models for each ROI and model using Stan. Across both ROIs, the peak of the posterior distributions of the estimated dimensionality for type II was higher than for type I, mirroring the structure of the two problems.

#### 4.3.3. Discussion

Besides knowing which areas show neural patterns with functional dimensionality, an important question concerns the degree of the underlying dimensionality. Using data from a categorization task where participants had to attend to either one or two features of a stimulus, we demonstrate how our method can be used to test whether the degree of underlying dimensionality of neural patterns varies with task demands. A notable strength of the dataset for our research question is that the authors used the same stimuli in a within-subject paradigm, counterbalancing the order of the two categorization tasks across subjects. This allowed us to investigate how the dimensionality of a neural pattern changes with task, while controlling for possible effects due to differences in signal-to-noise ratios across participants or brain regions.

Our results show that, as expected, the degree of underlying functional dimensionality is higher when the task required attending to two stimulus features instead of only one. Notably, this assumption was implicit to the conclusions drawn by the authors in the original publication (Mack et al., 2016). The authors analyzed neural patterns in hippocampus and implemented a cognitive model to show that stimulus-specific neural patterns were stretched across relevant compared to irrelevant dimensions. Thus, irrelevant dimensions were compressed and the dimensionality of the neural pattern was reduced the less dimensions were relevant to the categorization problem. Our approach allows to directly assess this effect without the need of fitting a cognitive model.

## 5. General Discussion

Multivariate and model-based analyses of fMRI data have deepened our understanding of the human brain and its representational spaces (Norman et al., 2006; Kriegeskorte and Kievit, 2013; Haxby et al., 2014; Turner et al., 2017). However, before evaluating specific representational accounts, it is sensible to first ask the more basic question of whether brain areas displays functional dimensionality more generally. Here, we presented a novel approach to estimate an area’s functional dimensionality by a combined SVD and cross-validation procedure. Our procedure identifies areas with significant functional dimensionality and provides an estimate, reflecting uncertainty, of the degree of underlying dimensionality. Across three different data sets, we confirmed and extended the findings from the original contributions.

After verifying the operation of the method with a synthetic (simulated) dataset in which the ground-truth dimensionalities were known, we applied our method to three published fMRI datasets. In each case, the procedure confirmed and extended the authors’ original findings, advancing our understanding of the function of the brain regions considered. Each of three datasets highlighted a potential use of estimating functional dimensionality.

In the first study, working with data from Mack et al. (2013), we demonstrated that testing for functional dimensionality can complement model-based fMRI analyses that evaluate more specific representational hypotheses. First, one cannot find a rich relationship between model representations and brain measures when there is no functional dimensionality in regions of interest. Second, there might be additional areas that display significant functional dimensionality that do not show correspondence with the model.

These additional areas invite further analysis as they might implement processes and representations outside the scope of the tested model. Functional dimensionality can indicate interesting unexplained signal. For example, in the first dataset examined, functional dimensionality was found in all the areas identified by Mack et al. (2013), plus medial BA 8, which is a candidate region for task difficulty and response conflict (see Alexander and Brown, 2011, for a model of medial prefrontal cortex function), which was not the authors’ original focus but may merit further study.

In the second study, working with data from Bracci and Op de Beeck (2016), we demonstrated how stimuli could be grouped or organized in different fashions to explore how dimensional organization varies across the brain. In this case, the data matrix was either organized along shape or category. We found neural patterns of shape and category selectivity consistent with the authors’ original results. However, we found the selectivity to be more mixed in our analyses and identified additional responsive regions, mirroring our results when we considered data from Mack et al. (2013).

Our method may have been more sensitive to signal because it makes fewer assumptions about the underlying representational structure and allows for individual differences in the underlying dimensions. In this sense, assessing functional complexity complements existing analysis procedures. Indeed, our approach could be used to evaluate multiple stimulus groupings to inform feature selection in encoding models (Diedrichsen and Kriegeskorte, 2017; Naselaris et al., 2011).

In a third study, working with data from Mack et al. (2016), we evaluated whether our method could identify changes in task-driven dimensionality. By combining estimates of functional dimensionality with a hierarchical Bayesian model, we found that the functional dimensionality in LOC was higher when a category decision required using two features rather than one. These results are consistent with the original authors’ theory but were hitherto untestable. In summary, assessing functional dimensionality across these three studies complemented the original analyses and revealed additional nuances in the data. In each case, our understanding of the neural function was further constrained. Moreover, comparing the results to those from model-based and other multivariate approaches was informative in terms of understanding underlying assumptions and their importance.

Of course, as touched upon in the Introduction, there are many possible ways to assess dimensional structure in brain measures and progress has been made on this challenge (Rigotti et al., 2013; Machens et al., 2010; Rigotti and Fusi, 2016; Diedrichsen et al., 2013; Bhandari et al., 2017; Lehky et al., 2014). Here, our aim was to specify a general, computational efficient, robust, and relatively simple and interpretable procedure that can easily be applied to whole brain data to first test for statistical significant functional dimensionality and, if found, to provide an estimate of its magnitude using Bayesian hierarchical modeling to make clear the uncertainty in that estimate.

We hope our contribution is useful to researches interested in further exploring their data, whether it be fMRI, MEG, EEG, or single-cell recordings. Researchers may consider variants of our method. For example, as mentioned in the Introduction, the SVD could be substituted with another procedure depending on the needs and assumptions of the researchers. There is no magic bullet to the difficult problems of estimating the underlying dimensionality of noisy neural data, but we have made progress on this issue both theoretically and practically. In doing so, we have also provided additional insights into the brain basis of visual categorization. We hope that by demonstrating the merits of estimating the functional dimensionality of neural data that we motivate others to take advantage of this additional and complementary viewpoint on neural function.

## 6. Data availability

A Matlab toolbox for estimating functional dimensionality of fMRI data as well as data needed to replicate the analyses presented here will be made available after publication. Nifti files and code for the analyses presented here are available from the authors upon request.

## 7. Acknowledgments

This work was funded by the National Institutes of Health [grant number 1P01HD080679]; the Leverhulme Trust [grant number RPG-2014-075]; and Wellcome Trust Senior Investigator Award [WT106931MA] to Bradley C. Love. Correspondences regarding this work can be sent to. The authors are grateful to all study authors for sharing their data and wish to thank all members of the LoveLab and Jan Balaguer for valuable input. Declarations of interest: none.

